# Antimicrobial Resistance Prediction in *Salmonella enterica* Using Frequency Chaos Game Representation and ResNet-18

**DOI:** 10.1101/2025.09.13.675356

**Authors:** Saifelden M. Ismail, Salma H. Fayed

**Affiliations:** Communication and Information Engineering Program, Zewail City of Science and Technology, Giza, Egypt

**Keywords:** Antimicrobial resistance, Deep learning, Frequency Chaos Game Representation (FCGR), ResNet-18, *Salmonella enterica*, *Staphylococcus aureus*, Genomic prediction

## Abstract

Antimicrobial resistance (AMR) prediction from bacterial genomes remains a major challenge for clinical microbiology and surveillance. We developed a deep learning model based on Frequency Chaos Game Representation (FCGR) and a ResNet-18 architecture to classify resistance phenotypes directly from whole-genome assemblies. Using homology-aware clustering to prevent genomic data leakage, we trained and evaluated models for *Salmonella enterica* (seven antibiotics) and *Staphylococcus aureus* (five antibiotics). The *Salmonella* model achieved high predictive accuracy, particularly for cephalosporins, while performance was lower for tetracycline and ampicillin. The *Staphylococcus aureus* model demonstrated that the pipeline generalizes to Gram-positive bacteria, with strong results for methicillin (Balanced Accuracy = 0.85). Benchmarking against the gene-based tool ResFinder showed that the FCGR-based model did not match the performance of ResFinder on most antibiotics, but achieved competitive results for cephalosporins. This study demonstrates the feasibility of applying FCGR-based deep learning to AMR prediction across bacterial species, though substantial improvements would be needed before clinical application.

## I. Introduction

Antimicrobial resistance (AMR) is one of the greatest health threats of the 21st century [1], causing around 1.2 million deaths each year with numbers continuing to rise [2]. Combined with the slow development of new antibiotics, this creates a pressing need for better ways to predict and monitor AMR.

Traditional antimicrobial susceptibility testing (AST) methods take 18 to 24 hours to identify bacteria and determine their susceptibility [2]. Because approximately 50% of antibiotic treatments begin without proper pathogen identification [3], there is an urgent need for faster and more accurate predictions of antimicrobial resistance.

Recent advances in artificial intelligence (AI) and machine learning (ML) have opened new opportunities to address these challenges. Deep learning methods have proven very effective in predicting AMR. Green et al. achieved 91.9% sensitivity for predicting resistance in *Mycobacterium tuberculosis* using convolutional neural networks (CNNs), which reduced diagnostic times from weeks to hours [4]. However, current ML-based AMR prediction methods have several drawbacks. A recent meta-analysis showed that while ML methods reach pooled area under the curve (AUC) scores around 0.82, they experience challenges with retrospective methodology, inconsistent data processing, and limited clinical validation [5].

Current genomic methods for predicting antimicrobial resistance primarily depend on databases that identify known resistance genes by matching patterns against established sources like ResFinder, NCBI, and CARD. Tools such as ABRicate [7] are the current standard for quick genotypic resistance prediction. However, they are limited because they rely on previously characterized resistance mechanisms. These methods cannot find new resistance patterns or reflect the complex, multi-gene nature of resistance that arises through various evolutionary pathways [8].

One promising way to tackle these limitations is by using alignment-free genomic representations. Chaos Game Representation (CGR) and its extension, Frequency Chaos Game Representation (FCGR), have become effective tools for transforming DNA sequences into high-density, information-rich 2D images. These images keep biological meaning while standardizing input sizes [9]. FCGR converts sequences of different lengths into equal-sized images based on k-mer frequencies [10]. This makes them ideal for deep learning applications. Recently, combining FCGR with CNNs for viral classification achieved 96.29% accuracy [11]. Ren et al. demonstrated that SNP encodings, including FCGR, along with machine learning methods, can provide strong predictive performance for antimicrobial resistance in *E. coli* [12]. However, their study only focused on predictions for one species, used limited evaluation metrics, and did not compare results against standard gene-based tools like ResFinder. Additionally, their data splitting method did not control for similarities between training and test isolates, raising concerns about genomic leakage.

Building on these foundations, this study improves alignment-free genomic AMR prediction by applying ResNet-18 architectures to FCGR of whole genome sequences. To ensure a thorough evaluation, we use homology-aware data splitting and compare our predictions against the gene-database-based tool ResFinder [15] as a relevant clinical baseline. To test cross-species generalizability, we also apply the same pipeline to *Staphylococcus aureus* (*S. aureus*), a clinically important Gram-positive pathogen. This work serves as a proof-of-concept to evaluate whether alignment-free, database-independent approaches can achieve meaningful predictive performance for AMR across bacterial species.

## II. Methods

### A. Dataset Description

The dataset used in this study was obtained from the Journal of Clinical Microbiology (JCM) publication [16], with project numbers PRJNA292661 and PRJNA292666. This dataset provided the foundation from which antibiotics of interest were selected with their corresponding samples. Only samples with labels for the chosen drugs were retained, while irrelevant samples were removed.

In addition to *Salmonella enterica*, we evaluated the proposed pipeline (shown in Figure 1) on a *Staphylococcus aureus* dataset compiled from two previously published collections [20], [21]. For drug selection, we followed a quantitative filtering procedure. First, we computed the percentage of missing labels (Null %), as drugs with extremely high missing values were less informative. Then, a score was calculated based on the resistant (R) to sensitive (S) ratio. The final combined score was calculated using the following formula, with thresholds for missing values (Null % *<* 50%) and minimal R/S ratio (*>* 5%):

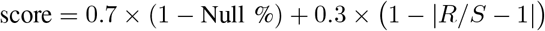

**Fig. 1.**
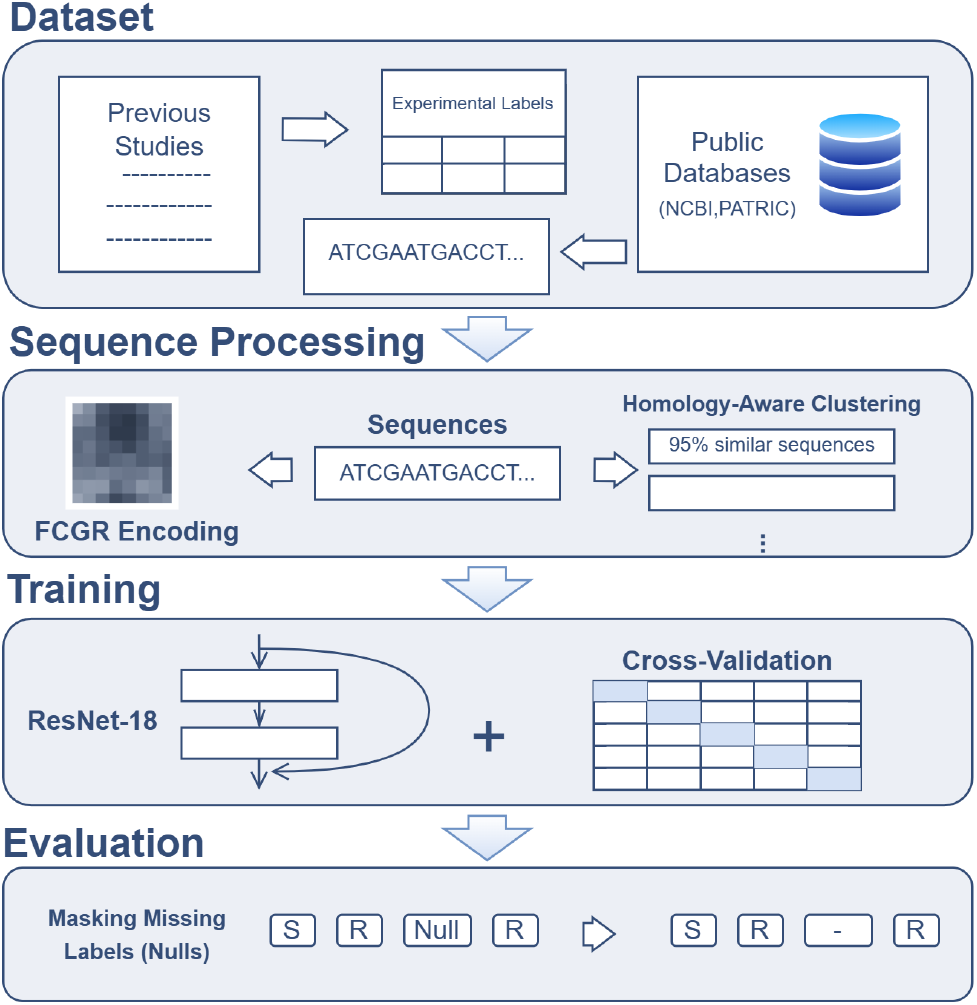
Workflow of the proposed pipeline.

Using this procedure, five antibiotics were selected for analysis: erythromycin, methicillin, ciprofloxacin, clindamycin, and penicillin. After filtering to retain only isolates with labels for at least one of these drugs, we obtained 5,883 genomes.

### B. Dataset Splits and Homology Considerations

Proper splitting was given significant focus in this study to prevent information leakage between homologous genomes and to ensure a fair evaluation of the model. Because of the large dataset size and computational constraints, utilizing standard homology-based clustering tools such as MMseqs2 [17] and CD-HIT [18] was not feasible. Instead, we employed a homology-aware clustering strategy using sourmash [19], which relies on MinHash sketching to approximate k-mer similarity.

For each dataset, genome assemblies were initially sketched with sourmash (version 4.8.6) using a k-mer size of 31 and a scaling factor of 1000, and pairwise similarities between all genomes were calculated with *sourmash compare* to obtain a distance matrix *d* = 1 − *s* (Jaccard similarity *s*). A threshold of *d* ≤ 0.05 (approximately 95% shared 31-mers, implying high average nucleotide identity when *k* = 31) was used to define closely related genomes. Smaller thresholds (e.g., *d* ≤ 0.01) would produce tighter, more homogeneous clusters, whereas larger thresholds (e.g., *d* ≤ 0.10) would yield clusters containing genomes with greater diversity; a threshold of *d* ≤ 0.05 maintained a balance between biological coherence and prevention of cluster leakage.

A graph was constructed in which genomes were represented as nodes, and edges were drawn between genomes with distances below the threshold (*d* ≤ 0.05). Connected components of this graph were then extracted using the networkx library, with each component representing a homology cluster. All genomes within a cluster were assigned entirely to either the training or the test set, and these clusters were used as grouping units for *GroupKFold* cross-validation to avoid any leakage between training and validation splits.

Because intact clusters vary in size, our target split naturally resolved to 12.43% test samples (632 genomes, 36 clusters) and 87.57% training samples (4452 genomes, 154 clusters) for *Salmonella*. The same procedure was applied to the *S. aureus* dataset, yielding 5,327 training samples (90.55%, 284 clusters) and 556 test samples (9.45%, 69 clusters). In both datasets, assigning intact clusters strictly to a single partition prevented leakage between the training pool and test set. Within the training pool, *GroupKFold* cross-validation further ensured that all genomes from the same cluster were assigned to the same fold, preventing any leakage between training and validation splits. Both datasets show class imbalance across labels, which is addressed by utilizing positive weights while training the model, as well as imbalance-aware metrics (e.g., balanced accuracy, MCC, Jaccard) during model evaluation. Class-specific positive weights were computed as 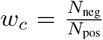, where *N*_pos_ and *N*_neg_ are the numbers of positive and negative samples for class *c* in the training set.

### C. Frequency Chaos Game Representation (FCGR)

Each genome assembly was converted into an FCGR matrix, which encodes the frequency of all possible *k*-mers in a two-dimensional grid [10]. We used the open-source *fcgr* Python library [22] to generate these matrices directly from FASTA/FNA files.

We evaluated multiple *k*-mer sizes (6, 7, and 8) during development of the *Salmonella* model, and based on empirical performance, we selected *k* = 8 for all subsequent experiments. The choice of *k* = 8 was further supported by biological reasoning: bacterial antimicrobial resistance genes are typically 500–1,500 bp in length, and an 8-mer representation generates 2^8^ *×* 2^8^ = 256 *×* 256 matrices comprising 65,536 unique *k*-mers, providing sub-gene-level resolution sufficient to capture conserved motifs within resistance determinants. For each genome, all contigs were concatenated into a single sequence prior to FCGR computation, and the resulting matrices were stored as .*npy* files to avoid repeated conversion during training. During training, each FCGR matrix was min– max normalized to the range [0, 1], preserving relative intensity patterns across genomes.

Figure 2 shows sample FCGR images for the two species included in this study. The FCGRs capture the k-mer frequency distribution of each genome. *S. aureus* exhibits a visually distinct, blockier structure compared to *Salmonella*, likely reflecting differences in GC content and genomic organization between Gram-positive and Gram-negative bacteria.

**Fig. 2.**
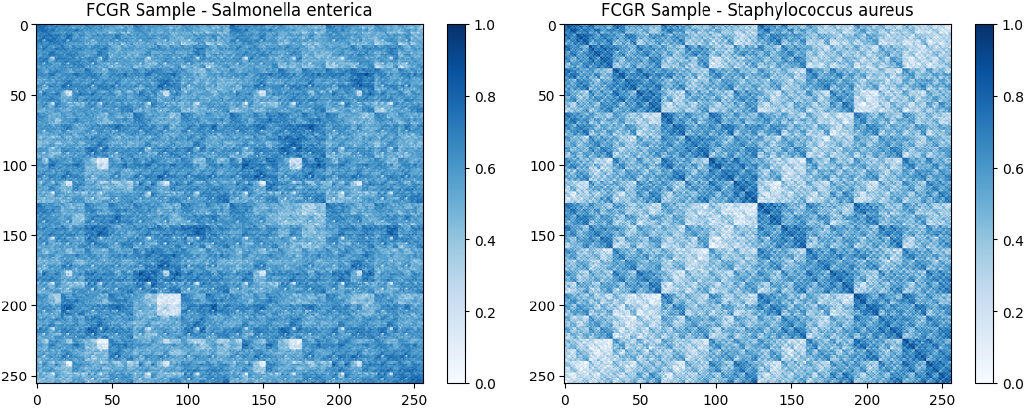
FCGR images of *Salmonella enterica* (left) and *Staphylococcus aureus* (right). Both 8-mer FCGR matrices were normalized and log-transformed to enhance contrast for display purposes. *S. aureus* exhibits a visually blockier pattern, reflecting its distinct genomic composition as a Gram-positive organism.

#### 1) Label Encoding and Preprocessing

Resistance phenotypes were binarized by encoding resistant (R) and intermediate (I) isolates as `0`, and susceptible (S) isolates as `1`. Although this representation is inverted compared to common conventions (where resistance is taken as the positive class), we adopted it because preliminary experiments indicated that the models converged more stably and achieved superior predictive performance when susceptibility was treated as the positive outcome. Intermediate isolates were merged with resistant due to their low frequency and biological plausibility. To ensure that this label choice did not bias evaluation, we report only symmetric, polarity-invariant metrics (Balanced Accuracy, MCC, and Jaccard), and handle missing labels (NULLs) by masking them on a per-drug basis during training and evaluation. Because susceptibility (S) is encoded as the positive class, all reported precision, recall, F1, and Jaccard values correspond to the susceptible class. Balanced Accuracy, MCC, and ROC AUC are polarity-invariant and remain unchanged regardless of which class is designated as positive.

### D. Deep Convolutional Neural Network Architecture

We employed a ResNet-18 architecture [23] as the backbone. ResNet-18 was selected due to its relatively small size, which is well-suited to the dataset scale, and because its residual connections help preserve weak but informative signals in the FCGR representations by allowing lower-level features to bypass layers and reach deeper layers unchanged. The first convolutional layer was modified to accept single-channel input, and the final fully connected layer was replaced with a linear classifier matching the number of antibiotic resistance classes. The model was trained from scratch without pretrained weights.

The *Salmonella* model was trained for 19 epochs and *S. aureus* model for 20 epochs, both with a batch size of Optimization was performed using the Adam optimizer (*β*_1_ = 0.9, *β*_2_ = 0.999, default values) with the one-cycle learning rate schedule as implemented in `fastai`. MixUp augmentation (mixing parameter *α* = 0.1) was applied as a regularization strategy. No early stopping was used; instead, model checkpoints were saved based on the best Matthews correlation coefficient (MCC) on the validation set, and convergence was assessed by monitoring validation metrics stabilization across epochs. The loss function was a masked, weighted binary cross-entropy, designed to account for class imbalance. Alternative imbalance-handling strategies such as oversampling, undersampling, and focal loss were not explored in this study but remain directions for future work. Evaluation was carried out using cluster-aware cross-validation with *GroupKFold* and clusters as the grouping variable.

## III. Results and Discussion

### A. Evaluation Metrics

We evaluated independent test set performance per drug (subsequently averaged) using standard `scikit-learn` metrics: balanced accuracy, precision, recall, F1, *F*_*β*_ (*β* = 2), ROC AUC, Jaccard index, and MCC, thresholding logits at 0.0. To align with the clinical goal of identifying viable treatments, Susceptible (S) was defined as the positive class (1) and Resistant (R) as negative (0); consequently, precision, recall (i.e., Sensitivity (S)), and F1-scores reflect the model’s ability to detect treatable isolates. Polarity-invariant metrics (MCC, balanced accuracy, Jaccard, ROC AUC) robustly handle class imbalance.

### B. Performance on Independent Test Set

On the independent test set, 6-mer, 7-mer, and 8-mer FCGR models achieved balanced accuracies of 0.84, 0.87, and 0.86 and MCC values of 0.56, 0.63, and 0.73, respectively; we therefore selected *k* = 8 for all subsequent experiments. Table I reports the final performance of the species-specific models on the independent test set using 8-mer FCGR images. For *Salmonella enterica*, the model demonstrated strong overall performance (Balanced Accuracy = 0.86, F1 = 0.94, MCC = 0.73) with near-perfect predictions for cefoxitin, ceftiofur, and ceftriaxone (Balanced Accuracy ≥ 0.94, F1 ≥ 0.99, MCC ≥ 0.84), but lower performance on tetracycline, ampicillin, and amoxicillin-clavulanic acid (Balanced Accuracy 0.71– 0.79, MCC 0.55–0.58).

**TABLE I:** Final performance of the species-specific FCGR + ResNet-18 models on the independent test set for 8-mer FCGR images. Metrics are reported overall and per drug, rounded to two decimal places.

| Species | Class | Bal. Acc | Precision | Recall | F1 | $F_\beta$ | ROC AUC | Jaccard | MCC |
| --- | --- | --- | --- | --- | --- | --- | --- | --- | --- |
| *Salmonella* | overall | 0.86 | 0.94 | 0.94 | 0.94 | 0.94 | 0.89 | 0.89 | 0.73 |
|  | tetracycline | 0.79 | 0.92 | 0.66 | 0.77 | 0.70 | 0.90 | 0.62 | 0.58 |
|  | ampicillin | 0.71 | 0.83 | 0.98 | 0.90 | 0.94 | 0.93 | 0.81 | 0.55 |
|  | amoxicillin-clavulanic acid | 0.74 | 0.92 | 0.97 | 0.95 | 0.96 | 0.87 | 0.90 | 0.57 |
|  | cefoxitin | 0.94 | 0.99 | 0.99 | 0.99 | 0.99 | 0.95 | 0.98 | 0.86 |
|  | ceftiofur | 0.98 | 1.00 | 0.99 | 0.99 | 0.99 | 0.99 | 0.99 | 0.92 |
|  | gentamicin | 0.86 | 0.97 | 0.99 | 0.98 | 0.99 | 0.97 | 0.96 | 0.80 |
|  | ceftriaxone | 0.96 | 1.00 | 0.98 | 0.99 | 0.98 | 0.98 | 0.98 | 0.84 |
| *S. aureus* | overall | 0.74 | 0.85 | 0.78 | 0.81 | 0.79 | 0.82 | 0.68 | 0.44 |
|  | erythromycin | 0.68 | 0.86 | 0.84 | 0.85 | 0.85 | 0.77 | 0.74 | 0.35 |
|  | methicillin | 0.85 | 0.99 | 0.73 | 0.84 | 0.77 | 0.96 | 0.72 | 0.64 |
|  | ciprofloxacin | 0.72 | 0.93 | 0.80 | 0.86 | 0.82 | 0.82 | 0.76 | 0.33 |
|  | clindamycin | 0.64 | 0.85 | 0.73 | 0.79 | 0.75 | 0.69 | 0.65 | 0.25 |
|  | penicillin | 0.83 | 0.61 | 0.82 | 0.70 | 0.76 | 0.89 | 0.54 | 0.60 |

The strong performance for cefoxitin, ceftiofur, and ceftriaxone can be explained by the correlation between these antibiotics. Because these cephalosporins share highly correlated resistance labels driven by common *β*-lactamase mechanisms, the multi-label setup effectively triples the available training signal for each drug. It is therefore possible that the model exploits shared label patterns rather than learning fully independent per-drug features. This interpretation is supported by the substantially lower performance on uncorrelated drugs such as tetracycline (Bal. Acc. = 0.79) and ampicillin (Bal. Acc. = 0.71), which cannot benefit from correlated labels.

Table II reports class-wise performance. Overall specificity was high (0.952) while sensitivity was lower (0.746), indicating resistant isolates are more likely to be missed. This is particularly evident for ampicillin (0.450) and amoxicillin-clavulanic acid (0.511), where resistant isolates are frequently misclassified as susceptible; by contrast, cefoxitin, ceftiofur, ceftriaxone, and tetracycline all have resistant sensitivity exceeding 0.88.

**TABLE II:** Sensitivity (Resistant) and Specificity (Susceptible) per antibiotic for *Salmonella enterica*.

| Class | Sensitivity (R) | Specificity (S) |
| --- | --- | --- |
| overall | 0.746 | 0.952 |
| tetracycline | 0.919 | 0.662 |
| ampicillin | 0.450 | 0.976 |
| amoxicillin-clavulanic acid | 0.511 | 0.972 |
| cefoxitin | 0.882 | 0.988 |
| ceftiofur | 0.978 | 0.990 |
| gentamicin | 0.730 | 0.993 |
| ceftriaxone | 0.950 | 0.980 |

### C. Staphylococcus aureus Performance

To assess whether the FCGR + ResNet-18 pipeline generalizes beyond Gram-negative pathogens, we trained a separate model on the *Staphylococcus aureus* dataset described in Section II. As summarized in Table I, the *S. aureus* model achieved a balanced accuracy of 0.74 and an MCC of 0.44 on the test set, with the strongest performance for methicillin (Bal. Acc. = 0.85, MCC = 0.64), consistent with the well-characterized *mecA*-mediated resistance mechanism, and solid performance for penicillin (Bal. Acc. = 0.83, MCC = 0.60) despite a heavily imbalanced label distribution (314 resistant vs. 93 susceptible in the test set). In contrast, erythromycin and clindamycin showed weaker balanced accuracy (0.68 and 0.64, respectively) and MCC values (0.35 and 0.25), likely reflecting the greater heterogeneity of macrolide and lincosamide resistance mechanisms in *S. aureus*. Class-wise, the model achieved a resistant sensitivity of 0.749 and susceptible specificity of 0.788, with methicillin showing both high sensitivity (0.979) and specificity (0.730), penicillin maintaining a balanced profile (0.843/0.817), and erythromycin trading higher specificity (0.843) for lower sensitivity (0.517), indicating a tendency toward false negatives that could lead to undertreatment. Compared with *Salmonella* (overall Bal. Acc. = 0.86, MCC = 0.73), the *S. aureus* model achieves more modest performance, as expected given the different drug panels, distinct Gram-positive genomic architecture, and potential differences in dataset quality, but nevertheless demonstrates that the FCGR-based pipeline is not restricted to a single species and can capture resistance-associated genomic patterns in a fundamentally different bacterial lineage.

### D. Cross-Validation Results

While the independent test set provides a strong indication of generalization ability, it represents only a single held-out split. To further assess the stability and robustness of model performance, we also conducted 5-fold cross-validation on the training pool. Across folds, the *Salmonella enterica* model achieved a mean balanced accuracy of 0.81 *±* 0.06 and an MCC of 0.58 *±* 0.10, with precision, recall, and F1 scores of around 0.90. Cross-validation of the *S. aureus* model yielded a mean balanced accuracy of 0.64 *±* 0.04 and an MCC of 0.28 *±* 0.06, confirming stable performance across folds, though with lower mean performance than the *Salmonella* model, reflecting the more heterogeneous drug panel. Detailed per-fold metrics are provided in the accompanying GitHub repository.

### E. Comparison to Baselines

To better contextualize the performance of our deep learning approach, we compared it against the widely used gene-based tool ResFinder. We evaluated predictions from ABRicate [7] using the ResFinder resistance gene database [15]. For each genome assembly, ABRicate was run across all samples, and the resulting resistance gene labels were mapped to antibiotic classes using a curated drug–class dictionary. ResFinder evaluation for *Staphylococcus aureus* followed the same methodology as for *Salmonella enterica*: ABRicate was run with the ResFinder database to detect resistance genes, and the resulting drug-class labels from the `raw_classes` column were mapped to phenotypic antibiotic columns using the same curated dictionary. Samples were predicted as resistant (R) if any corresponding resistance gene was detected and susceptible (S) otherwise, and were then evaluated against ground-truth phenotypic labels using the same eight metrics (balanced accuracy, precision, recall, F1, F_*β*_, ROC AUC, Jaccard, and MCC) with resistance as the positive class.

When comparing the ResFinder results with the *Salmonella enterica* model, several important contrasts emerge. For tetracycline, ResFinder reports a high BA of 0.98 and MCC of 0.96, whereas the CNN model achieves only 0.79 BA and 0.58 MCC. For ampicillin, ResFinder shows strong results (BA = 0.96, MCC = 0.94) in contrast to the CNN model (0.71 BA, 0.55 MCC). In contrast, for cefoxitin, ResFinder achieves a BA of 0.97 and MCC of 0.88, but the CNN model performs competitively with a BA of 0.94 and MCC of 0.86. The remaining cephalosporins (ceftiofur and ceftriaxone) achieve BA values of 0.98 and 0.96 and MCC values of 0.92 and 0.84, respectively, comparable to the best performances reported in ResFinder. Taken together, ResFinder consistently outperforms the CNN model across a majority of drugs. The CNN model narrows the gap for cephalosporins (cefoxitin, ceftiofur, ceftriaxone), but underperforms substantially on tetracycline, ampicillin, and amoxicillin-clavulanic acid. These results indicate that while FCGR-based deep learning can achieve reasonable predictive performance for some drug classes, it does not currently offer advantages over established gene-based methods like ResFinder. Full per-drug ResFinder metrics are provided in the online repository.

The results for *S. aureus* follow the same trend across all evaluated drugs: erythromycin (BA = 0.90 vs. 0.68, MCC = 0.79 vs. 0.35), clindamycin (BA = 0.87 vs. 0.64, MCC = 0.74 vs. 0.25), and penicillin (BA = 0.88 vs. 0.83, MCC = 0.73 vs. 0.6), demonstrating that gene-based methods remain superior for this species despite the CNN model’s competitive performance on methicillin. McNemar’s test showed statistically significant differences (*p <* 0.05) between CNN and ResFinder predictions for all evaluated drugs in both species.

To further benchmark against alternative approaches, we trained several additional models on the same *Salmonella* dataset. A Random Forest classifier using gene presence/absence features achieved modest performance, with balanced accuracy of 0.56–0.65 across drugs and MCC values of 0.14–0.51, indicating poor discrimination of resistant isolates.

We also implemented a simple 3-block CNN comprising three convolutional layers with batch normalization and ReLU activations, interleaved with pooling layers, followed by an adaptive average pooling and a fully connected classifier. It yielded moderate performance (overall balanced accuracy 0.69, masked MCC 0.26), indicating limited ability to capture subtle patterns in the FCGR representations. Larger architectures achieved better results; EfficientNet reached overall balanced accuracy 0.85 (ranged 0.71–0.95 across drugs) and MCC 0.58 (0.46–0.69). Overall, these results indicate that deeper CNN architectures can better detect FCGR features and approach, but do not surpass, the performance of ABRicate with ResFinder.

### F. Model Interpretability

To investigate which sequence features the model attends to, we generated saliency maps for 16 ceftiofur-resistant *Salmonella* genomes and used them to rank *k*-mers by their contribution to the model output. Across genomes, the union of the top salient *k*-mers yielded 2,814 globally important *k*-mers, of which 102 (3.6%) fell within the clinically relevant *β*-lactamase *blaCMY-2*, accounting for 5.0% of the total saliency mass. This limited overlap indicates that the model’s predictions are not primarily driven by *blaCMY-2* alone but by *k*-mers distributed more broadly across the genome, which may reflect lineage effects or population structure—features that correlate with resistance phenotypes due to clonal inheritance rather than causal mechanisms. Without phylogenetic controls or experimental validation, these saliency maps should therefore be interpreted as a descriptive analysis of model behavior rather than evidence of novel biological insights.

### G. Limitations and Future Work

While our results demonstrate the feasibility of FCGR-based deep learning models for predicting antimicrobial resistance, several important limitations must be acknowledged. First, although training the ResNet-18 models was manageable, generating FCGR matrices was computationally intensive and limited exploration of deeper architectures and extensive hyperparameter searches; we did not perform formal ablations for MixUp, class weighting, thresholding, nor alternative imbalance-handling strategies. Second, FCGR encodes only *k*-mer frequencies, discarding positional context (e.g., plasmid vs. chromosomal origin); future work could explore ordered FCGR variants to retain this. Third, despite evaluating two species (*S. enterica* and *S. aureus*), broader multi-species validation across additional pathogens is needed to establish generalizability. Finally, several barriers remain for clinical deployment, including regulatory requirements for in vitro diagnostics (IVD), the need for interpretable models that clinicians can trust, computational constraints in microbiology laboratories, and the necessity of prospective clinical validation; closing the performance gap with gene-based methods like ResFinder is a key prerequisite. Future work will focus on addressing these limitations by collecting larger, fully labeled multi-species datasets, conducting formal ablations and architecture comparisons, and incorporating phylogenetic controls to distinguish genuine resistance-associated signals from confounding population structure.

## IV. Conclusion

In this work, we evaluated FCGR paired with ResNet-18 for predicting antimicrobial resistance directly from bacterial genomes, without predefined resistance gene databases. The *Salmonella enterica* model achieved high predictive accuracy for cephalosporins but underperformed on antibiotics such as tetracycline and ampicillin, while the *Staphylococcus aureus* model demonstrated that the approach generalizes to Gram-positive bacteria, with particularly strong results for methicillin (Bal. Acc. = 0.85, MCC = 0.64). Compared with the gene-based tool ResFinder, our model did not match its performance on most antibiotics, and ResFinder remains more accurate for the drugs tested, but homology-aware clustering ensured that these comparisons were not confounded by genomic leakage. Curated datasets are often small and lack complete labels, and preprocessing pipelines for FCGR require significant computing power; future work should focus on creating larger, high-quality multi-species datasets, exploring alternative architectures, and conducting formal ablation studies to close the performance gap with curated gene-based methods and move toward systems that can be used in clinical settings.

## Data Availability

All code, dataset description files, and generated tables are available at https://github.com/XHCFS/amr-fgcr-prediction-project; raw sequencing data are accessible from the public repositories cited in the Methods.

## Acknowledgment

We thank Sama S. Eltaher (Zewail City University of Science and Technology) for biological insights and guidance on bioinformatics tools, and Mariam I. Mohamed for assistance with data collection.

